# Host availability, climate and pupal-stage ecology shape the predicted environmental suitability of the New World screwworm across Central and South America

**DOI:** 10.64898/2026.11.05.764564

**Authors:** Lucie Blasco, Marina Voinson, Miguel Moreno, Danilo Escobar, Marine Combe, Rodolphe E. Gozlan

**Author notes:** Corresponding author: Rodolphe E. Gozlan.

## Abstract

**Background:** The New World screwworm, *Cochliomyia hominivorax*, causes primary myiasis in warm-blooded vertebrates. Its recent resurgence in Central America has renewed the need for spatial tools that identify environmentally suitable areas and surveillance gaps. Most previous distribution models have emphasized climate and livestock, whereas ecological constraints acting during the free-living pupal stage have rarely been incorporated.

**Methods:** We compiled 2,461 georeferenced occurrence reports from the literature across Central and South America and linked them to climate, livestock density, forest cover and superficial soil predictors. Four sequential (nested) logistic models quantified the incremental contribution of predictor blocks, compared using likelihood-ratio tests, AIC and discrimination metrics. Logistic regression and Random Forest were then compared using five-fold spatial block cross-validation, which we treat as the primary measure of generalisation, and an independent municipality-level 70:30 train-test split. Variable importance was assessed using Random Forest Mean Decrease Accuracy together with partial-dependence profiles. Because presence and pseudo-absence classes were balanced before fitting, model outputs are interpreted as relative environmental suitability rather than absolute probability of occurrence.

**Results:** The literature database represented 2,589 infected hosts, dominated by cattle, pigs, dogs and humans. Across the sequential models, AUC increased from 0.780 for climate alone to 0.898 with livestock density, 0.908 with forest cover and 0.918 with superficial soil properties; the increment attributable to the soil block (ΔAUC = 0.010) was smaller than the between-fold variability observed under spatial cross-validation (SD 0.049) and is therefore reported as suggestive rather than established. Sheep and horse density and cold-quarter soil temperature showed the highest Mean Decrease Accuracy values, while silt texture ranked above cattle density and forest cover. Under spatial block cross-validation Random Forest was markedly more stable than the GLM (mean AUC 0.926 ± 0.049 versus 0.814 ± 0.157) and was retained as the final model. On the independent municipality-level test set it achieved AUC 0.981, TSS 0.824, sensitivity 0.850 and specificity 0.972, values we regard as optimistic relative to the spatially blocked estimates. Predicted suitability was highest across Central America and parts of northern and tropical South America.

**Conclusions:** *C. hominivorax* occurrence was associated with interacting climatic, host and landscape conditions. Superficial soil variables added a small, biologically interpretable signal consistent with constraints acting during pupation, but one that lies within the uncertainty of our spatial validation and requires confirmation with independent data. The maps estimate relative environmental suitability rather than infection incidence, and are intended to identify areas where surveillance could be strengthened rather than to guide control operations directly.

## Background

Parasitic diseases emerge from complex interactions among hosts, environmental conditions and human activities, making eco-epidemiological approaches particularly relevant when part of the parasite life cycle occurs outside the host (Francesconi & Lupi, 2012; Ung et al., 2021; Deiana et al., 2024). The New World screwworm, *Cochliomyia hominivorax* (Coquerel, 1858), is a particularly important example. This calliphorid fly is an obligate parasite whose larvae develop in living tissues and cause primary myiasis in livestock, wildlife and humans. The species has major veterinary and economic consequences in the Americas, with cattle production receiving especially intensive attention (Costa-Júnior et al., 2019).

The life cycle of *C. hominivorax* spans ecological compartments that are likely to impose different constraints on persistence. Females oviposit on wounds or mucosal surfaces, larvae develop in living tissues, and mature larvae leave the host and enter the soil, where pupation takes place. The duration and success of this free-living stage are influenced by environmental conditions including temperature, precipitation, water drainage and other physico-chemical properties of superficial soils (Fresia et al., 2021; Akhoundi et al., 2023). Adult stages then emerge and disperse, over long distances of 20 km or more when hosts are scarce (Valdez-Espinoza et al., 2025). Consequently, the geographical niche of the species should reflect not only climate and host availability, but also ecological constraints acting during pupation.

Historically, *C. hominivorax* occurred from the United States through Central America and into South America. Large-scale sterile insect programmes progressively eliminated the fly from North and Central America and established a biological barrier in Panama, while the species remained endemic in parts of South America (Fresia et al., 2013). The introduction into Libya and subsequent eradication also illustrate the capacity of the species to establish outside its native range and the importance of coordinated control (Cunningham et al., 1992; Lindquist et al., 1992). Recent re-emergence in Central America has renewed major concerns over its capacity for geographical expansion and highlighted the need for surveillance tools that can identify favourable environments even where reported cases remain sparse (Keatts et al., 2025; Valdez-Espinoza et al., 2025). This management history carries a direct methodological consequence: for several decades the fly was absent from large areas of Central America that remained environmentally suitable, so historical absence in that region reflects successful intervention rather than environmental unsuitability. Any distribution model assembled from records spanning the eradication era must therefore be interpreted with the footprint of those programmes in mind.

Previous ecological niche analyses have largely emphasised climatic predictors and, in some cases, livestock density (Mulieri & Patitucci, 2019; Valdez-Espinoza et al., 2025). Complementary work in the Caribbean has combined environmental correlates of occurrence with genetic analyses to investigate the species’ geographical distribution and population differentiation (Tietjen et al., 2022). This leaves a biologically important gap, since the free-living pupal stage occurs in soil. Yet, edaphic variables are rarely incorporated explicitly into continental-scale distribution models. In parallel, the reported host spectrum is broad but unevenly documented, making it difficult to distinguish ecological host importance from surveillance intensity.

Here, we developed a large-scale predictive model of *C. hominivorax* occurrence in Central and South America using a literature-derived occurrence database and environmental predictors representing climate, domestic host availability, forest cover and superficial soil properties. We asked whether progressively incorporating these biologically distinct predictor groups improved model performance, and whether a non-linear Random Forest model generalized more consistently in space than logistic regression. We further used the compiled literature to describe the reported host spectrum and mapped environmentally suitable areas with few nearby documented cases. We hypothesised that the geographical distribution of *C. hominivorax* reflects interacting constraints acting across multiple stages of its life cycle, and specifically that soil variables relevant to pupation provide information beyond climate and host availability.

## Methods

### Literature search and study selection

A literature search was conducted in Google Scholar and Web of Science between April and May 2026 and covered publications available up to 8 May 2026, without language restriction. The geographical scope comprised Belize, Costa Rica, El Salvador, Honduras, Panama, Nicaragua, Guatemala, Brazil, Colombia, Argentina, Venezuela, Guyana, French Guiana, Uruguay, Peru, Chile, Paraguay, Bolivia, Ecuador, Suriname and the Falkland Islands/Malvinas. The search combined “Cochliomyia hominivorax” with country names; the complete search strings for each database are given in Supplementary Material (Table S1).

Records were eligible if they reported one or more confirmed cases of *C. hominivorax* myiasis in any host, in any of the countries listed above, with a location that could be georeferenced at or below country level. Reviews without primary case data, records identifying only *Cochliomyia* to genus level, and laboratory or colony studies without field cases were excluded. The search returned 12,002 references, of which 7,349 duplicates were removed, leaving 4,653 records for title and keyword screening (Fig. 1). Screening excluded 3,991 records falling outside the study area or subject, leaving 662; a further 203 duplicate or off-topic records, 161 records without data meeting the database parameters, and 195 records without clinical cases or consisting of bibliographic studies were excluded at successive full-text stages. In total, 103 publications met the eligibility criteria and yielded the 2,461 georeferenced case reports analysed here. Records were screened in Rayyan (Ouzzani et al., 2016; Harrison et al., 2020). Because full-text PDFs could not be automatically associated with the screened Rayyan records, NotebookLM was used as an assistance tool during full-text screening and data extraction, an application that has recently been evaluated for ecological systematic reviews (Costas et al., 2026). Specifically, it was used to screen full-text articles against the eligibility criteria above and subsequently to identify case reports among those retained. All final inclusion and exclusion decisions and all extracted information were manually checked against the source documents. Reporting follows the PRISMA 2020 statement where applicable (Page et al., 2021); the review protocol was not prospectively registered.

**Figure 1.**
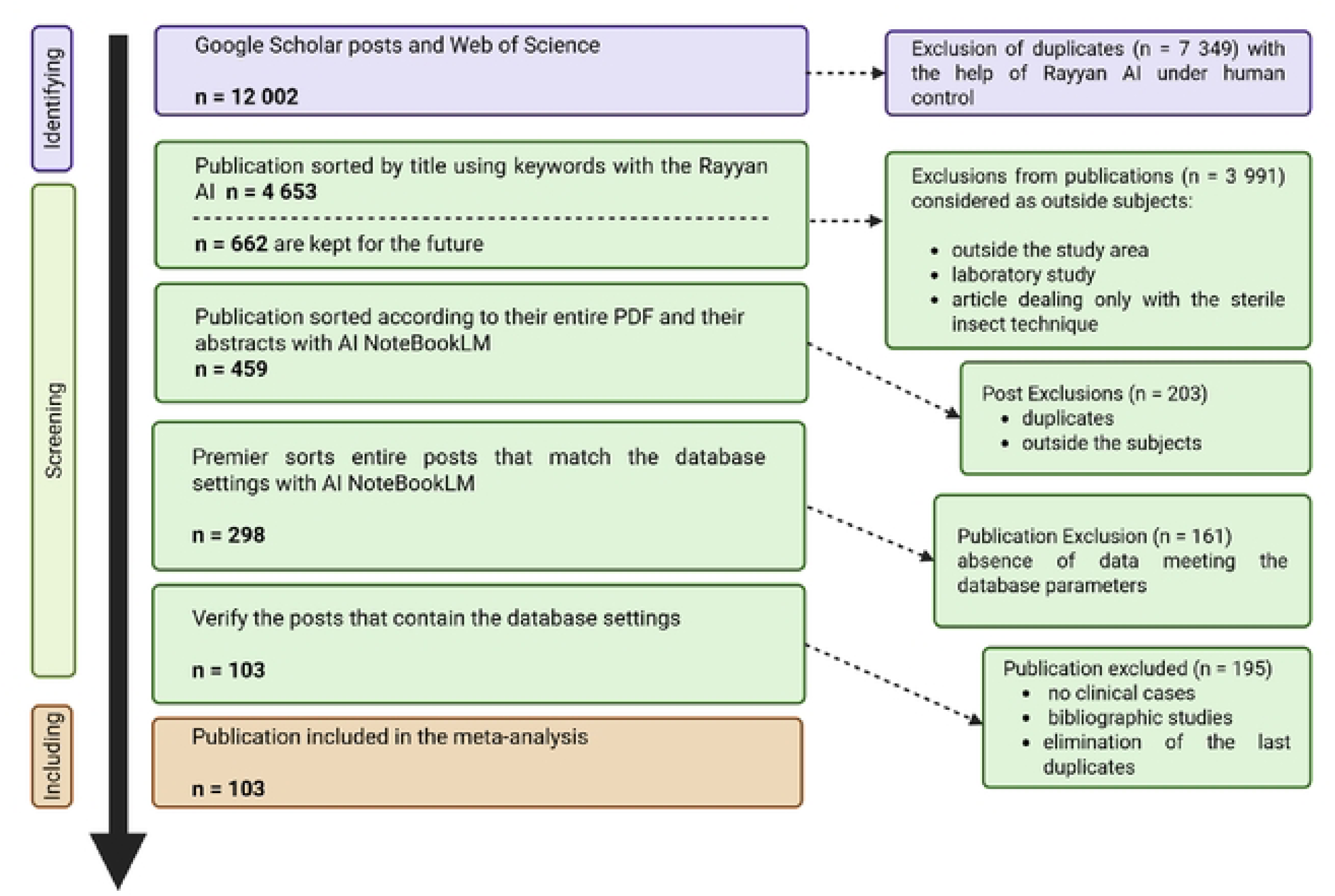
PRISMA 2020 flow diagram of literature identification, screening and inclusion used to build the occurrence database (see Methods).

### Data extraction and georeferencing

For eligible records, we extracted country, province or region, municipality, geographical coordinates when reported, host species, host age and sex where available for all host types including humans, total number of cases, reported comorbidities or pre-existing conditions for all host types, sampling dates, information on the likely place of infection, and data type. For domestic animals we recorded the location of the farm where the case was diagnosed, which can differ from the location of infection where animals are moved between pastures; for dogs and cats the location of the veterinary clinic was used where no other information was available; for human cases we used the place of residence when recorded, the city of infection for travel-related cases, or, failing both, the address of the admitting hospital. Where no location could be established below country level, coordinates were assigned to the geographic centre of the country (n=12, 0.49% of cases). Missing coordinates were otherwise assigned from reported municipalities using Google Earth and cross-checked. Locations displayed in maps from Rodríguez-Hidalgo et al. (2018) and Maxwell et al. (2015) were digitised in QGIS version 4.0.2. The year of each reported case was recorded where available, and the temporal distribution of records is summarised in Supplementary Material (Fig. S1).

### Environmental predictors

Predictors were selected a priori according to the biology of *C. hominivorax* and represented ecological constraints acting on adult, host-associated and pupal stages. Air temperature and precipitation variables were obtained from WorldClim 2 at approximately 1-km resolution, representing the 1970-2000 climatology (Fick & Hijmans, 2017). Soil moisture information was derived from Trabucco and Zomer (2019). Mean conditions and variables describing the warmest/wettest and coldest/driest quarters were considered in order to capture seasonal extremes across the study region. Soil temperature data were obtained from Lembrechts et al. (2022), who provide gridded estimates for the 0-5 cm and 5-15 cm layers; we used the 0-5 cm layer as the closest available approximation to the conditions experienced during shallow pupation. Livestock density layers were derived from the Gridded Livestock of the World database for the 2010 reference year (Gilbert et al., 2018). Superficial soil properties included drainage, available water capacity and percentages of clay, sand, gravel and silt, based on the WISE30sec harmonised soil property values (Batjes, 2016). Forest cover and forest-loss information were obtained from Hansen et al. (2013). The source, reference year, native resolution and transformation applied to each predictor are summarised in Supplementary Material (Table S2). Because the climate layers are long-term climatologies and the livestock and forest layers correspond to fixed reference years, whereas occurrence records span several decades, all predictors describe long-term average environmental conditions rather than conditions at the time of each individual report. Environmental values were extracted at occurrence and pseudo-absence locations in QGIS version 4.0.2.

### Pseudo-absences and data preparation

Pseudo-absence locations were generated randomly in QGIS outside a 20-km exclusion buffer around presence locations, a distance chosen to reflect the approximate dispersal range reported for adult screwworm flies. Random background sampling does not reproduce the uneven sampling effort of the occurrence data, so part of the discrimination reported below may separate surveyed from unsurveyed areas rather than suitable from unsuitable ones; this issue, and the sensitivity of the results to the buffer distance, are addressed in the Limitations. After environmental extraction, 2,461 presence locations and 2,363 pseudo-absence locations had usable predictor data. Random resampling was used to balance the two classes, giving a final modelling dataset of n = 4,922 records. Because class prevalence was therefore set by design rather than observed, fitted values are relative suitability scores and not calibrated probabilities of occurrence. Non-finite values were recoded as missing. Quantitative predictors used for modelling were transformed as sign(x) × log(|x| + 1), preserving the sign of negative temperature values while reducing skewness; all coefficients and odds ratios reported below are consequently expressed per unit of the transformed predictor rather than per native unit.

### Predictor selection and nested models

Spearman correlations were calculated among quantitative predictors because predictor distributions were non-normal, and the full correlation matrix is given in Supplementary Material (Fig. S3). Highly redundant variables were removed with reference to |r| ≈ 0.75 and to biological relevance. Where candidate climatic variables remained strongly correlated, simplified and fuller climate blocks were compared empirically. To quantify the incremental contribution of ecological predictor groups, four sequential logistic models were fitted to the same complete-case dataset: model A included climate variables; model B added domestic livestock densities; model C added forest cover; and model D added superficial soil properties. Because these models are nested, they were compared using likelihood-ratio tests and AIC in addition to AUC, TSS and LogLoss, so that the contribution of each block could be assessed against sampling variability rather than by change in discrimination alone.

### GLM and Random Forest modelling

Two complementary modelling approaches were compared. Logistic regression (GLM) was used to estimate the direction and magnitude of associations, expressed as odds ratios per unit of the signed-log-transformed predictors, whereas Random Forest was used to accommodate non-linear relationships and interactions. Random Forest variable importance was assessed using Mean Decrease Accuracy and interpreted alongside partial-dependence profiles. Model performance was assessed using AUC, TSS and LogLoss. Generalisation was evaluated using five-fold spatial block cross-validation, which reduces the inflation of performance caused by geographical proximity between training and validation records, and we treat this as the primary criterion for model selection. Spatial blocks were defined as a regular grid with block size determined automatically by blockCV and using repearted random assignment over 50 iterations. The final models were additionally evaluated on an independent 70:30 train-test split in which each municipality, and every point within it, was assigned exclusively to either the training or the test set, so that geographically coincident records could not appear in both.

Because neighbouring municipalities remain spatially autocorrelated at the continental scale, this split is less stringent than spatial blocking and its results are reported as a secondary and probably optimistic estimate. Sensitivity and specificity were calculated at the threshold that maximised TSS (0.43).

### Spatial prediction and surveillance-gap mapping

The selected Random Forest model was projected across the study region after harmonising predictor extent and resolution to a common grid and applying the same signed-log transformation used for model fitting. Forest-cover data were aggregated from their native high resolution to the common 1-km prediction grid. A calibration-domain layer was used to flag areas requiring environmental extrapolation, and is presented in Supplementary Material (Fig. S2) so that predictions made outside the calibration envelope can be identified. Projected values are mapped as relative environmental suitability rather than as probabilities of occurrence. A second map combined predicted suitability with the distribution of literature-derived cases, distinguishing favourable locations with and without a reported case within 20 km. Because suitability is itself estimated from those same occurrence records, this overlay is partly self-referential, and the locations it identifies are interpreted as hypotheses about surveillance effort rather than as confirmed high-incidence areas.

### Statistical analysis

Statistical analyses were conducted in R version 4.5.1 using dplyr (v1.2.1), tidyr, ggplot2 (v4.0.3), corrplot, randomForest (v4.7-1.2), patchwork (v1.3.2), iml (v0.11.4), sf (v1.1.1) and gt (v1.3.0), together with pROC (v1.18.5) for AUC computation, terra (v1.9.27) for raster manipulation and for spatial block cross-validation blockCV (v3.2-0). Spatial data preparation and mapping were performed in QGIS version 4.0.2. Analysis code and the derived occurrence database are available as described under Availability of data and materials.

## Results

### Incremental contribution of ecological predictor groups

Annual temperature, annual precipitation and annual soil temperature were removed in favour of seasonal variables because they were strongly correlated with their quarterly counterparts (r = 0.91, 0.96 and 0.95, respectively; Supplementary Material, Fig. S3). Although warmest- and coldest-quarter air temperatures were also correlated (r = 0.85), retaining the fuller climate block improved AUC from 0.75 to 0.78 and it was therefore retained. Across the sequential GLMs, discrimination improved as biologically distinct predictor groups were added: AUC increased from 0.780 for climate alone to 0.898 after adding livestock density, 0.908 after adding forest cover and 0.918 after adding soil properties, while TSS increased from 0.310 to 0.688 across the same sequence (Table 1). The increments attributable to forest cover and to soil properties (ΔAUC = 0.010 each) are an order of magnitude smaller than that attributable to livestock density (ΔAUC = 0.118), and smaller than the standard deviation of AUC across spatial validation folds (0.049); the corresponding likelihood-ratio tests and AIC values are reported in Table 1. Removal of sheep and goat density reduced the full-model AUC from 0.918 to 0.897and available water capacity contributed little additional discrimination (AUC 0.916 without versus 0.918 with the variable). The latter was removed from the final model for parsimony, a decision taken after inspecting model performance and therefore likely to render the final figures slightly optimistic.

**Table 1.** Incremental performance of sequential logistic models incorporating climate, livestock density, forest cover and superficial soil variables. ΔAUC is relative to the preceding model.

| Modele | AUC | TSS | LogLoss | delta_AUC | AIC | delta_AIC | LRT_chi2 | LRT_df | LRT_p |
|---|---|---|---|---|---|---|---|---|---|
| A – Climate | 0.7796 | 0.3103 | 0.5811 | NA | 5624.97 | NA | NA | NA | NA |
| B – Climate + Hosts | 0.8973 | 0.6645 | 0.3918 | 0.1177 | 3807.36 | -1817.61 | 1827.6 | 5 | 0 |
| C – Climate + Hosts + Forest | 0.9069 | 0.6719 | 0.3787 | 9,6 × 10<sup>-3</sup> | 3682.63 | -124.73 | 126.73 | 1 | 2,1233 × 10<sup>-29</sup> |
| D – Climate + Hosts + Forest + Soil | 0.9176 | 0.6881 | 0.3604 | 1,07 × 10<sup>-2</sup> | 3516.35 | -166.28 | 176.28 | 5 | 3,3300 × 10<sup>-36</sup> |

### Variable importance and direction of associations

In the Random Forest, the highest Mean Decrease Accuracy values were observed for sheep density (72.65), horse density (68.79) and cold-quarter soil temperature (51.17). Silt texture was the highest-ranked soil variable (36.11), ranking above cattle density (32.01) and forest cover (30.83) (Fig. 2). The GLM clarified the direction of associations; all odds ratios are expressed per unit of the signed-log-transformed predictor. Horse density (OR = 2.12) and pig density (OR = 1.79) were positively associated with occurrence (both p < 0.001), whereas cattle, goat and sheep densities showed negative coefficients (OR = 0.69, 0.29 and 0.70, respectively; all p < 0.001). Warm-quarter air temperature (OR = 4.70, p = 0.005), warm-quarter soil temperature (OR = 2.41, p < 0.001), wettest-quarter precipitation (OR = 1.77, p < 0.001), driest-quarter precipitation (OR = 1.10, p = 0.032) and cold-quarter soil temperature (OR = 1.70, p = 0.003) were positively associated with occurrence. Cold-quarter air temperature was not significant (p = 0.94). Forest cover was negatively associated with occurrence (OR = 0.78, p < 0.001). Full coefficients with 95% confidence intervals are given in Supplementary Material (Table S3). The combination of a high Random Forest importance for sheep density with a negative GLM coefficient indicates that this variable does not act as a simple index of host availability. Within the study region high sheep densities are concentrated in Patagonia and the high Andes, so the variable may partly encode geographical position and its associated climate. Because correlated predictors and geographically structured livestock systems complicate causal interpretation, all coefficients are treated as model associations rather than as direct mechanistic effects.

**Figure 2.**
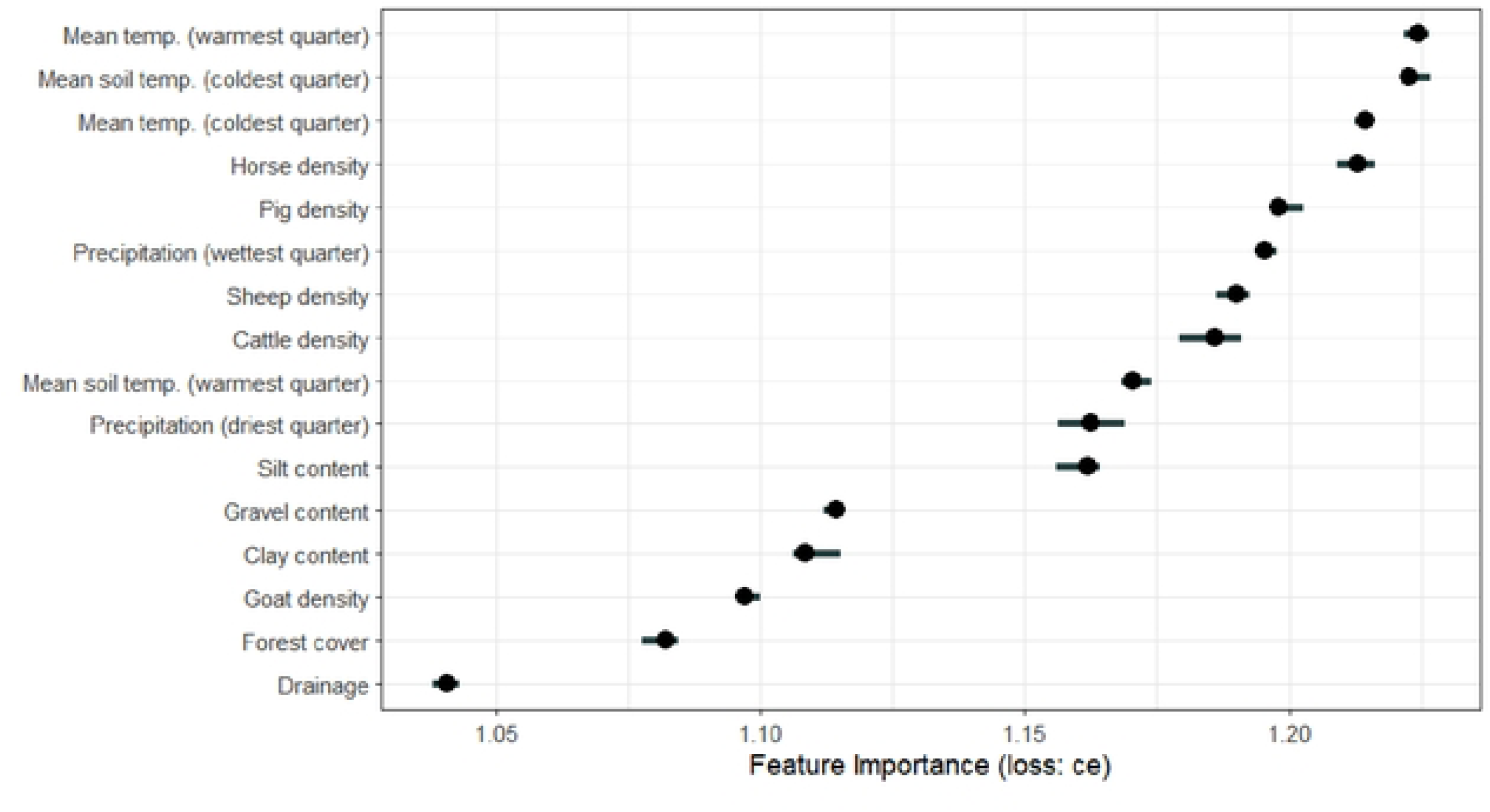
Random Forest variable importance (Mean Decrease Accuracy) for the final model.

### Spatial validation and model selection

Spatial block cross-validation provided our primary comparison between the two approaches. Random Forest retained a mean AUC of 0.926 (SD 0.049; range 0.87-0.99), compared with 0.814 (SD 0.157; range 0.57-0.95) for the GLM, and mean TSS was 0.67 and 0.53, respectively (Table 2, Fig. S3). Random Forest was therefore selected as the final predictive model because of its greater stability among geographical blocks. On the training data Random Forest showed near-perfect fit (AUC = 1.000, TSS = 1.000, LogLoss = 0.043), whereas the GLM achieved AUC = 0.916, TSS = 0.691 and LogLoss = 0.362; these training figures are reported for completeness and are not evidence of predictive performance. On the independent municipality-level 30% test set, Random Forest achieved AUC = 0.981, TSS = 0.824, LogLoss = 0.221, sensitivity = 0.850 and specificity = 0.972, and the corresponding GLM values were AUC = 0.892, TSS = 0.644, LogLoss = 0.403, sensitivity = 0.781 and specificity = 0.862. That these test values exceed the spatially blocked estimates is expected, because municipality-level partitioning does not remove spatial autocorrelation between neighbouring units; the blocked estimates are the more conservative description of expected performance in new areas.

**Table 2.** Comparative performance of GLM and Random Forest under five-fold spatial block cross-validation (the primary selection criterion) and on the independent municipality-level 70:30 test set. Spatial block values are mean ± SD across folds.

| Model | Evaluation | AUC | TSS | LogLoss | Sensitivity | Specificity |
|---|---|---|---|---|---|---|
| Random Forest | Spatial block CV | 0.926 ± 0.049 | 0.67 | — | — | — |
| GLM | Spatial block CV | 0.814 ± 0.157 | 0.53 | — | — | — |
| Random Forest | Independent test | 0.981 | 0.824 | 0.221 | 0.850 | 0.972 |
| GLM | Independent test | 0.892 | 0.644 | 0.403 | 0.781 | 0.862 |

### Predicted geographical suitability and surveillance gaps

Predicted relative suitability ranged from 0 to 0.968 and was spatially heterogeneous (Fig. 3). The highest predicted suitability occurred across Central America and north-western South America, with additional areas exceeding 0.95 in Ecuador, Peru, Colombia, Brazil, Venezuela and Uruguay. Lower suitability was predicted in the southernmost parts of the continent, particularly Chile and southern Argentina. Areas for which one or more environmental layers were unavailable are masked, and areas falling outside the calibration envelope are flagged in Supplementary Material (Fig. S2). The surveillance-gap overlay identified favourable areas lacking a literature-derived case within 20 km, including sectors of Colombia and northern Venezuela along the Andes, parts of Peru, the Amazon estuary and areas south of Buenos Aires (Fig. 4). In Central America, broad areas were predicted to be favourable while remaining incompletely represented in the literature-derived occurrence database, including parts of El Salvador.

**Figure 3.**
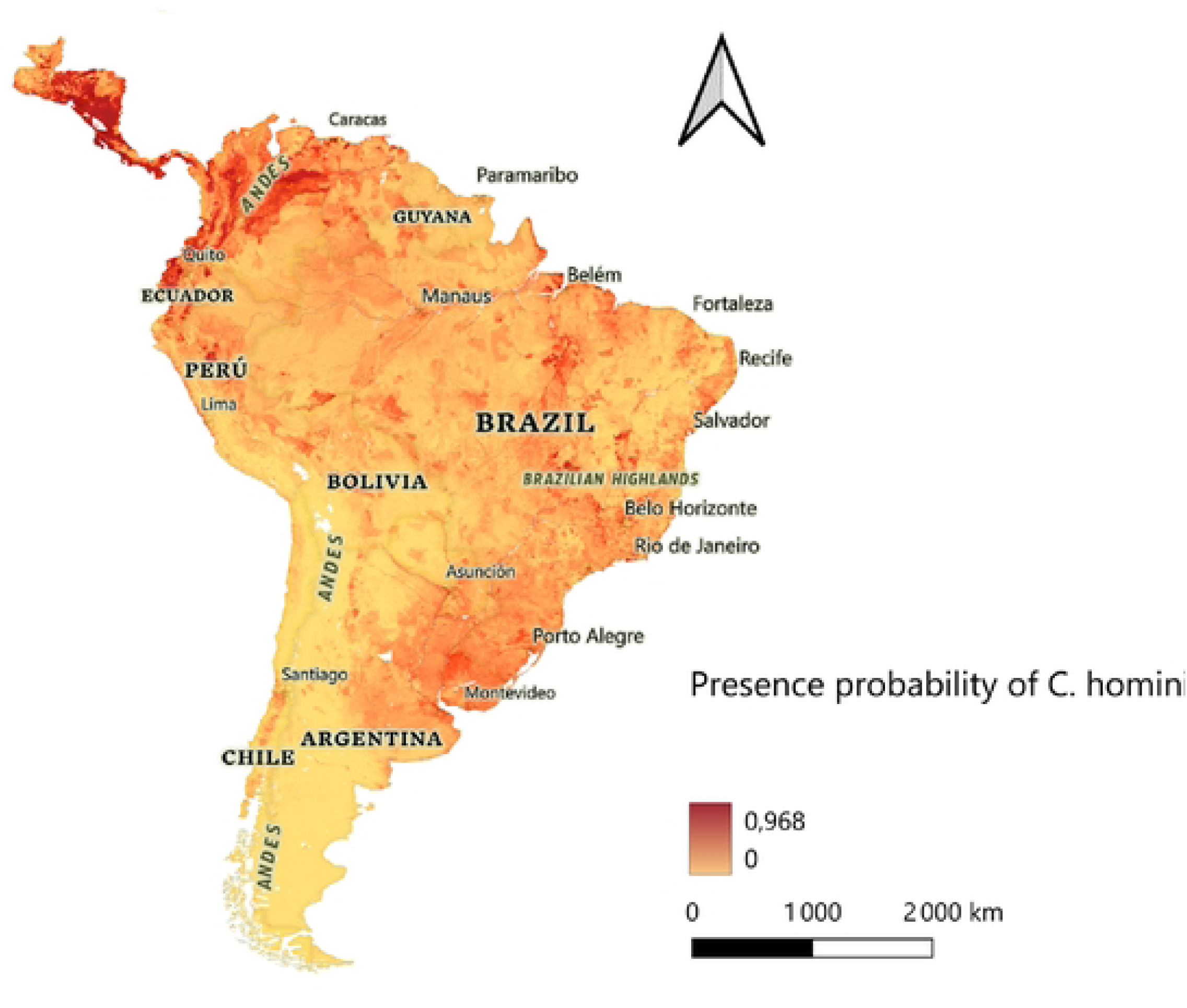
Predicted relative environmental suitability for *Cochliomyia hominivorax* across the study region from the final Random Forest model. Values are relative suitability scores derived from a class-balanced model and are not calibrated probabilities of occurrence.

**Figure 4.**
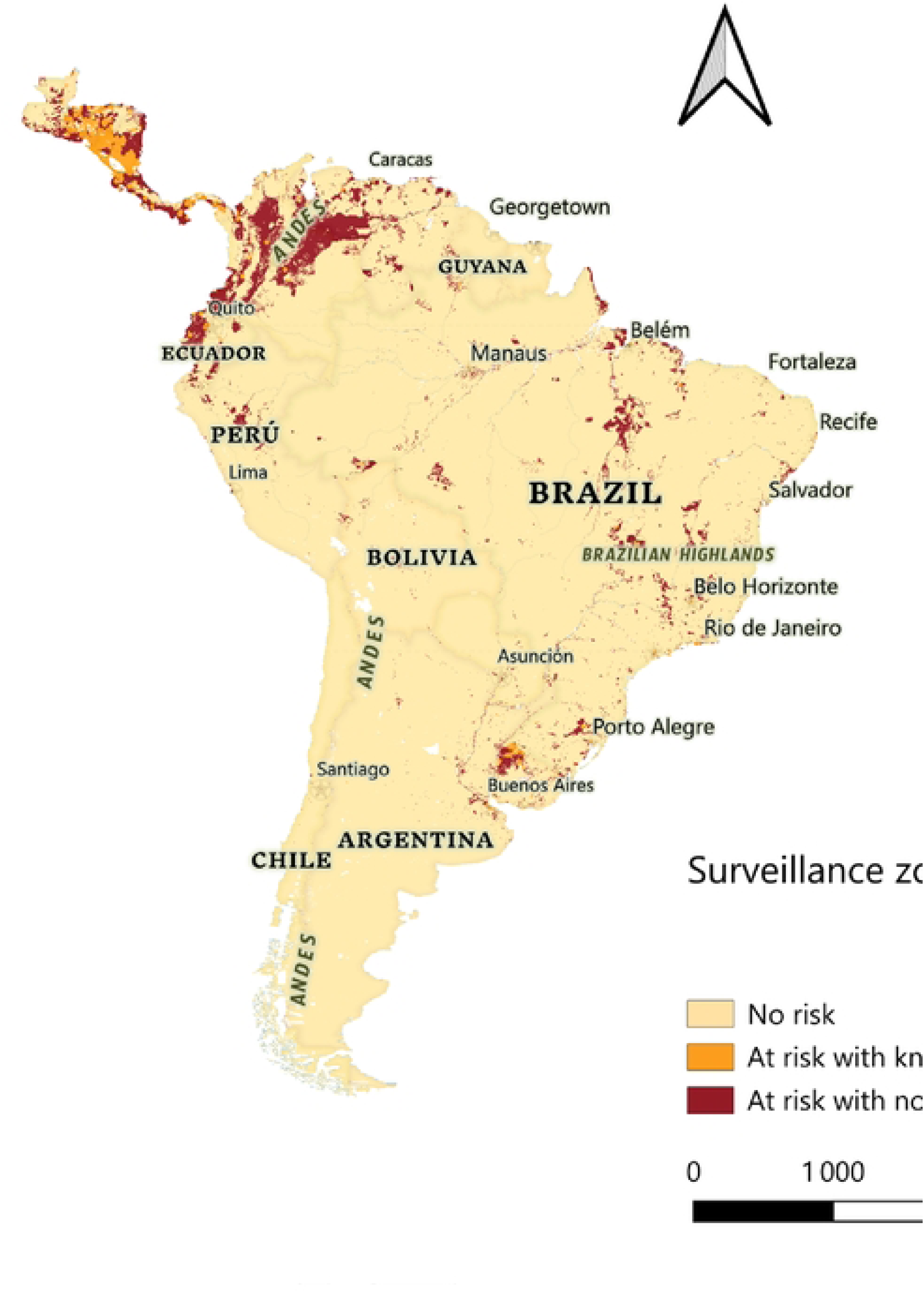
Surveillance-gap map combining predicted suitability with literature-derived occurrences. High-suitability areas lacking a reported case within 20 km are candidate surveillance priorities, not confirmed high-risk areas.

### Reported host spectrum

The database contained 2,461 case reports representing 2,589 infected hosts, because some reports included more than one host (Fig. 5). Cattle accounted for 58.8% of reported hosts, followed by pigs (11.8%), dogs (8.8%) and humans (6.5%). Wildlife represented only 0.6% of reported hosts but included taxonomically diverse mammals and birds, among them otters, cervids, rodents and egrets. Seasonal patterns differed among host groups: cattle, pigs and horses showed a peak in June, whereas goats and sheep peaked in November (Fig. 6). Dates were available for n = 1779 records, and missing dates limited inference for cats, wildlife and humans. Open wounds and traumatic lesions were the principal reported entry sites. Among the n = 153 human reports that included clinical detail, the associated conditions most frequently described were chronic disease, reduced capacity for self-care and neurological disorders; most reported human cases occurred between approximately 50 and 75 years of age, and the majority were male. All of these descriptive frequencies reflect what has been published rather than what has occurred, and should not be interpreted as unbiased estimates of host-specific infection risk or of patient characteristics.

**Figure 5.**
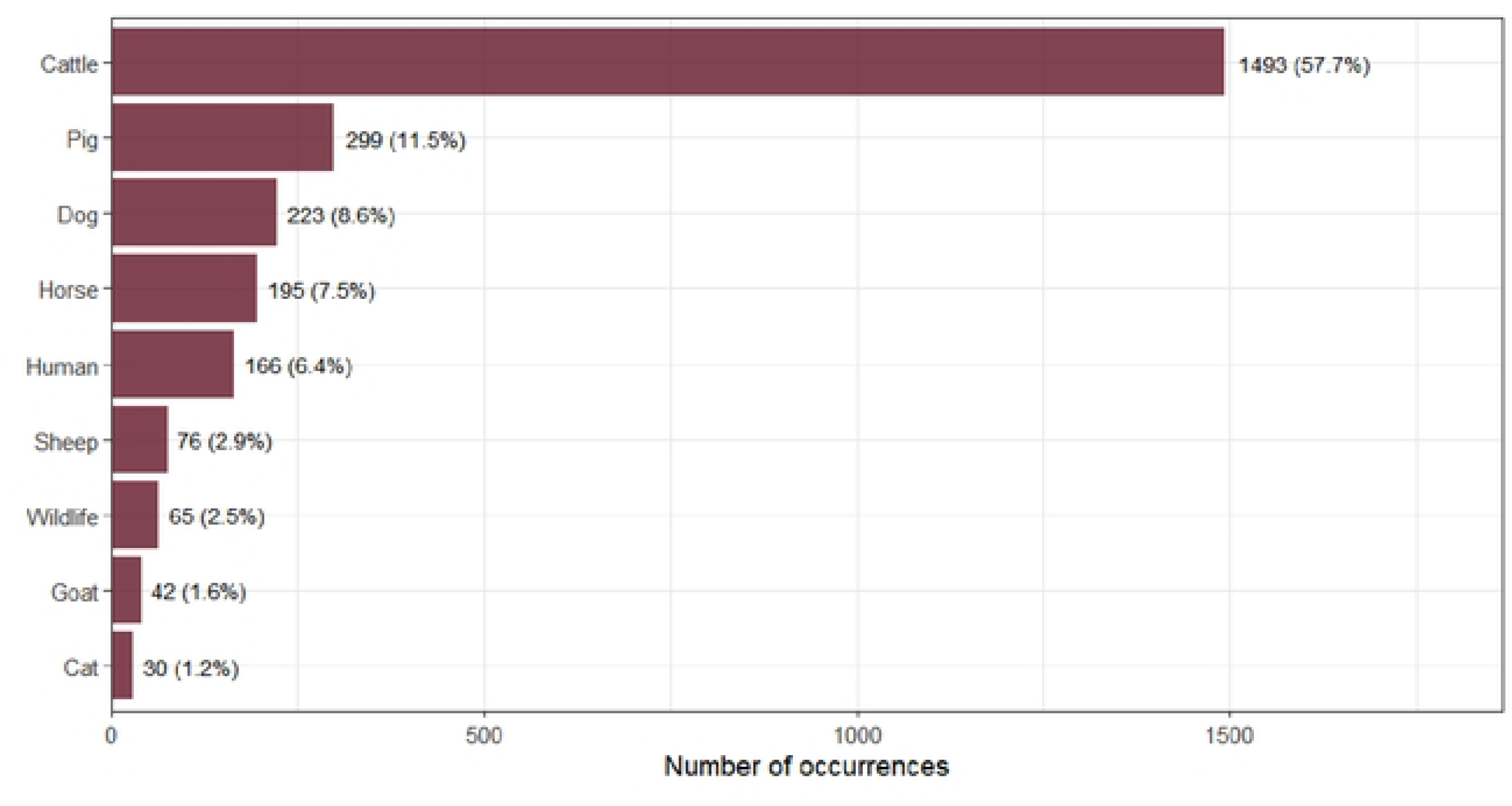
Distribution of reported host categories in the literature-derived database. Frequencies describe the published record and should not be interpreted as unbiased host-specific infection risk.

**Figure 6.**
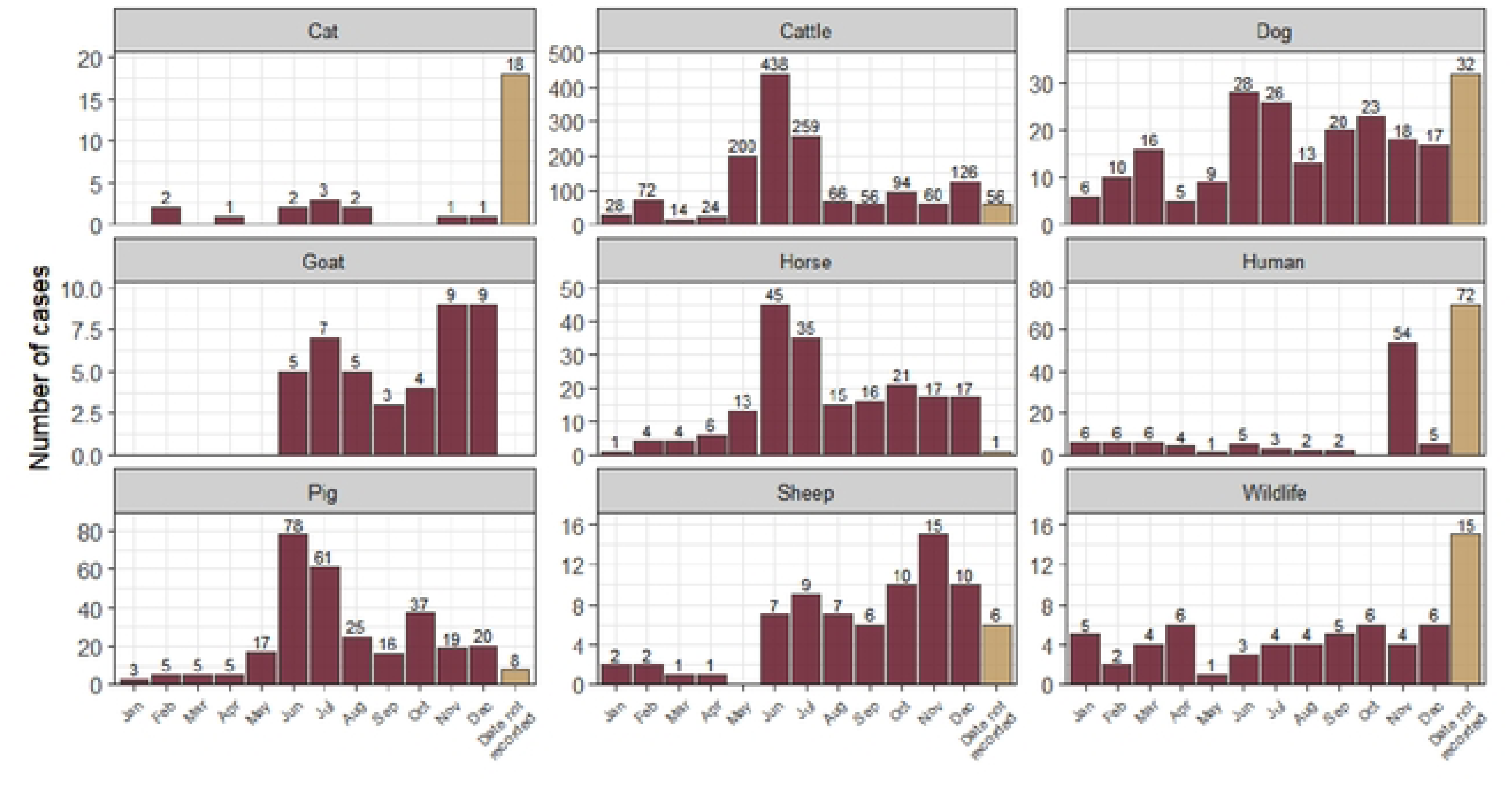
Distribution of cases by host type and time of year.

## Discussion

This study supports a multi-factor view of the ecological niche of *C. hominivorax*. Climate and domestic-host variables accounted for most of the gain in predictive performance, but forest cover and superficial soil properties provided additional discrimination. The latter result is biologically important because the pupal stage occurs outside the host and is directly exposed to soil temperature, moisture and texture (Oliveira, 1980). Distribution models that focus exclusively on adult climate envelopes or host availability may therefore omit constraints acting during such critical free-living stage (Mulieri & Patitucci, 2019; Valdez-Espinoza et al., 2025). The contribution of soil predictors was modest in absolute terms and should not be overstated. The ΔAUC associated with the soil block was 0.010, which is smaller than the variation in AUC across spatial validation folds, so on the present data this improvement is suggestive rather than demonstrated. Nevertheless, silt texture ranked above cattle density and forest cover in Random Forest importance, and cold-quarter soil temperature was among the three most influential predictors, patterns consistent with the hypothesis that pupation conditions act as an ecological filter. The distinction between statistical improvement and biological interpretation matters here. The present analysis does not demonstrate a causal effect of soil properties on pupal survival; it identifies a stage-specific environmental signal that merits targeted field and laboratory validation, and that should be tested in a design with greater power to separate edaphic from climatic variation (Fresia et al., 2021).

Livestock availability was the largest source of improvement beyond climate, but the ranking of livestock predictors requires cautious interpretation. Sheep and horse densities were more important in the Random Forest than cattle density, even though cattle dominated the published host records (Costa-Júnior et al., 2019). This mismatch likely reflects, at least in part, differences between ecological information and surveillance effort, and in part the geographical structure of livestock systems. As noted above, sheep density in this region largely delineates Patagonia and the high Andes and may therefore act as a spatial rather than an ecological covariate. Cattle myiasis has substantial economic consequences (Grisi et al., 2014) and is therefore intensively reported, whereas livestock-density layers can also capture broader farming systems, landscape structure and correlated environmental conditions (Costa-Júnior et al., 2019; Phillips et al., 2004). Differences between Random Forest importance rankings and GLM coefficients are not unexpected, because the two approaches quantify different aspects of predictor effects. The GLM estimates average conditional associations, whereas Random Forest importance reflects the contribution of a variable to prediction, including non-linear relationships and interactions with other environmental variables (Elith & Leathwick, 2009). Positive GLM associations for horses and pigs and negative coefficients for cattle, goats and sheep should consequently not be interpreted as host preferences. Future models should explicitly separate host abundance, husbandry practice, wound prevalence and surveillance intensity, and should test whether livestock predictors retain importance once geographical position is controlled for. Field observations in Venezuela highlight wounds associated with calving, castration, branding and dehorning as relevant contexts for infestation, reinforcing the need to consider husbandry practices alongside host abundance (Coronado & Kowalski, 2009). Seasonal patterns in case reporting differed among host groups, with cattle, pigs and horses peaking in June and goats and sheep in November. This difference may partly reflect distinct husbandry practices rather than biological differences in host susceptibility. In Andean regions of South America, small ruminants are commonly subject to seasonal transhumance between lowland and highland pastures (Rodríguez-Hidalgo et al. 2018), which could expose these animals to favourable transmission zones at different times of the year.

Forest cover was negatively associated with occurrence, suggesting higher predicted suitability in open or anthropogenic landscapes. This pattern may reflect the concentration of livestock and human activity in cleared landscapes rather than a direct inhibitory effect of forest itself (Francesconi & Lupi, 2012). Interfaces between recently deforested areas, pasture, wildlife and domestic animals could create conditions that simultaneously increase host availability and opportunities for parasite movement (Keatts et al., 2025; Voinson et al., 2026). The predicted high-suitability areas in parts of the Amazon basin therefore deserve particular caution. They may represent genuine livestock-wildlife interfaces, but they also include regions where occurrence data are sparse and where the model extrapolates beyond its calibration envelope (Fig. S2).

Spatial validation was central to model selection. The near-perfect training performance of Random Forest indicates substantial capacity to fit the calibration data and, considered alone, could suggest overfitting (Elith & Leathwick, 2009). However, performance remained high and comparatively stable under geographical block cross-validation, whereas the GLM varied markedly among spatial blocks. For a model intended to support surveillance across a large and environmentally heterogeneous region, stability across geographical partitions is more informative than training-set discrimination. The independent test results are consistent with the selected model, although municipality-level partitioning is a weaker guarantee of independence than spatial blocking, and external validation with newly collected surveillance data would provide a stronger test than any partition of the present database. We also note that we assessed discrimination only; the calibration of predicted suitability against observed occurrence frequency was not evaluated, and a presence-background calibration measure such as the continuous Boyce index would be a useful addition.

The surveillance-gap map is best interpreted as a hypothesis-generating tool. High predicted suitability in the absence of a nearby literature-derived case does not demonstrate undetected transmission, because absence from the database may result from low surveillance, non-publication, imperfect georeferencing or true absence. A review of human myiasis in Colombia similarly reported a concentration of published cases in regions with greater access to healthcare and surveillance infrastructure, illustrating how reporting patterns can shape the apparent geographical distribution of disease (Donato-Rondón et al., 2025). Nevertheless, these mismatches can help prioritise locations for active veterinary, wildlife and environmental surveillance. The literature-derived host synthesis reinforces this point. Wildlife accounted for very few published records despite the broad host range of *C. hominivorax* (Bautista-Garfias et al., 2023), suggesting that sylvatic infections are particularly under-characterised (Timbie et al., 2026). Feral swine and mobile bird species have been proposed as potentially informative sentinels or transport hosts, but their epidemiological role requires direct study (Altuna et al., 2021; Timbie et al., 2026).

An important distinction is therefore required between predictive contribution and mechanistic inference.

### Limitations and future directions

Several limitations qualify these results. First, occurrence records were derived from heterogeneous published sources and are spatially biased towards better-studied areas (Elith & Leathwick, 2009), whereas pseudo-absences were drawn at random across the study region; part of the discrimination we report may therefore separate surveyed from unsurveyed areas rather than suitable from unsuitable ones. Repeating the analysis with a target-group background, constructed from the localities of other field studies in the same literature, would test how far this affects the results. Second, the 20-km exclusion buffer used to generate pseudo-absences was not formally tested for sensitivity, and alternative buffer distances may produce different pseudo-absence distributions. Third, presences were not spatially thinned, so clusters arising from intensively documented areas, including the locations digitised from published maps, contribute repeated information; the number of distinct 1-km grid cells occupied is a more conservative measure of sample size than the number of reports. Fourth, records for which no sub-national location was available were assigned to country centroids (0,49% of all records), which can place presences in environments where the species may not occur. Fifth, in some human reports hospital location was used when the exposure site was unavailable, potentially shifting records towards urban centres. Finally, environmental predictors are long-term climatologies or fixed-year layers, whereas records span several decades, so the model describes average rather than contemporaneous conditions.

These limitations also define a clear route towards a One Health surveillance framework. Spatial suitability models can be combined with veterinary case reporting, wildlife surveillance and targeted field sampling rather than used as stand-alone control maps (Hailu et al., 2025). Species-level identification remains essential, because myiasis surveillance can detect other causative flies during and following C. hominivorax eradication programmes (Bermúdez et al., 2007). Detection of *C. hominivorax* DNA from environmental samples such as air and swabs is an intriguing possibility for detecting the fly in farm environments, but it remains prospective and requires assay development and field validation before operational use. Likewise, the present maps should not be used directly to determine sterile-insect release intensity; such deployment requires population-density estimates, reinvasion dynamics and regional operational planning. Population-genetic evidence provides complementary information on population structure and demographic history that cannot be inferred from environmental suitability maps alone (da Silva e Souza et al., 2025).

## Conclusion

Large-scale occurrence of *C. hominivorax* is best explained by a combination of climatic conditions, domestic-host availability and landscape characteristics. Superficial soil variables provided additional information consistent with ecological constraints during pupation, although the magnitude of that contribution remains within the uncertainty of our spatial validation and requires confirmation. A Random Forest model showed high and comparatively stable discrimination under spatial block validation and identified broad areas of environmental suitability across Central America and tropical South America. The principal contribution of this study is therefore not a claim that soil dominates screwworm distribution, but evidence that the ecology of a neglected free-living life stage is worth incorporating into spatial prediction, together with an explicit account of what literature-derived models of this kind can and cannot support. Used cautiously, the model provides a framework for targeting field surveillance and for integrating animal, human and environmental information in the context of New World screwworm re-emergence.

## Author contributions

LB compiled and curated the literature-derived database, conducted the spatial and statistical analyses. REG conceived and supervised the study and wrote the manuscript. MV, MC, DE and MM, reviewed and discussed the interpretation of the results.

